# Antarctica’s wilderness has declined to the exclusion of biodiversity

**DOI:** 10.1101/527010

**Authors:** Rachel I. Leihy, Bernard W.T. Coetzee, Fraser Morgan, Ben Raymond, Justine D. Shaw, Aleks Terauds, Steven L. Chown

## Abstract

Recent assessments of the biodiversity value of Earth’s dwindling wilderness areas^1,2^ have emphasized the whole of Antarctica as a crucial wilderness in need of urgent protection^3^. Whole-of-continent designations for Antarctic conservation remain controversial, however, because of widespread human impacts and frequently used provisions in Antarctic law for the designation of specially protected areas to conserve wilderness values, species and ecosystems^4,5^. Here we investigate the extent to which Antarctica’s wilderness encompasses its biodiversity. We assembled a comprehensive record of human activity on the continent (~ 2.7 million localities) and used it to identify unvisited areas ≥ 10 000 km^2^ ^(1,6-8)^ (i.e. Antarctica’s wilderness areas) and their representation of biodiversity. We show that, at best, 7 770 000 km^2^ of wilderness remains, covering 56.9% of the continent’s surface area, however it captures few important biodiversity features. Important Bird Areas^9^, ice-free Antarctic Conservation Biogeographic Regions crucial for biodiversity^10,11^, and areas with verified biodiversity records^12^ are largely excluded. Our results demonstrate that Antarctica’s wilderness has already declined to the exclusion of much of its biodiversity. But that on a continent set aside as a natural reserve^13^, increased regulation of human activity and urgent expansion of the Antarctic specially protected area network could feasibly reverse this trend.

---

Wilderness areas are important in the Earth system for maintaining biodiversity and large-scale ecosystem services^1,14^. They also provide baselines for assessing current and future anthropogenic environmental impacts elsewhere^15^. Catastrophic declines in Earth’s wilderness^2^ have led to urgent calls for action to secure it, including through the establishment of comprehensive targets^3^. The protection of Antarctica has been identified as crucial to this action^3^. Antarctica plays a significant role in the global climate system^16^, has unusual and surprisingly extensive biodiversity^10^, and despite its isolation, is under growing human pressure^16,17^. The actual area of wilderness on the continent remains unknown, as does the extent to which it captures Antarctic biodiversity, which is largely restricted to ice-free areas^11^. Arguments have been made that the whole of Antarctica can be considered, by default, a wilderness with the highest global level of conservation protection^18,19^. Provisions in the Protocol on Environmental Protection to the Antarctic Treaty for specially protected areas ‘…to protect outstanding environmental, scientific, historic, aesthetic or wilderness values…’ (Annex V to the Protocol), along with the growing scope of human impacts^20,21^ demonstrate, however, that this is not the case^5,8,22^.

We determined the area of wilderness on the continent, the extent to which it encompasses the continent’s ice-free, relatively biodiversity-rich ecoregions (known as Antarctic Conservation Biogeographic Regions (ACBRs)^12^), and the degree to which Important Bird Areas (IBAs)^9^, Antarctic Specially Protected Areas (ASPAs) and terrestrial biodiversity are captured within it. We also examined the spatial distribution of wilderness with respect to known threats to Antarctic biodiversity, specifically human activity^21,22^, non-native species^23^ and climate change^11^. Elsewhere, globally-significant wilderness areas are defined as those ≥ 10 000 km^2^ (1 million hectares)^1^, mostly intact (≥ 70% of historical habitat extent), and with low human population densities (≤ 5 people per km^2^)^1^. Because Antarctica has no industrial, urban or agricultural land-use^24^, Antarctic wilderness areas are often defined as those with no evidence of human activity^6,7,19^. Antarctic wilderness areas have, however, also been defined as those with an absence of human infrastructure or those with either no, or non-transient, human activity^8^. Here, we define an Antarctic wilderness area as any contiguous land area ≥ 10 000 km^2^, with no evidence of historical human visitation. Antarctica’s wilderness is the sum of these areas. We have adopted a stringent definition since human activities in Antarctica, even if transitory, can have large impacts, especially because of the slow biological rates of the indigenous terrestrial biota^25,26^, and because those activities are diversifying and growing rapidly on the continent^20-22^.

We assembled an extensive record of ground-based human activity across the Antarctic continent and its immediate offshore islands, from publications and scientific databases, spanning 1819 to 2018 (~ 2.7 million activity localities; Extended Data Fig. 1) (see ‘Data availability’) (hereafter, the primary data). These human activity data were reported or recorded in a variety of formats, necessitating protocols to ensure compatibility for use on a 5 × 5 km equal area grid to define areas of visitation (Methods). To identify Antarctica’s wilderness areas, the high-resolution (25 km^2^) activity grid was aggregated to a 2 500 km^2^ resolution grid (excluding marine cells). Any four contiguous cells (i.e. ≥ 10 000 km^2^) with no activity data were defined as a wilderness area. To determine whether Antarctica’s wilderness captures major features of biodiversity value, biodiversity spatial layers were used. These are Antarctica’s ACBRs, IBAs, species locality records captured by the Antarctic Terrestrial Biodiversity Database, and the ASPAs^9,12^. In advance of this assessment, biological sampling records were excluded from the primary data, and a constrained wilderness identified, to avoid biasing the analysis because most biodiversity records come from human visits to areas. The biodiversity layers were overlaid onto this constrained wilderness and representation calculated as a proportion of overlap. Current threats to wilderness areas were evaluated as proximity to scientific and tourism activity^21,22^, establishment suitability for non-native species^23^, and expected changes in ice coverage^11^, using the wilderness grid derived from the primary data. Because ice-free areas contain the majority of Antarctica’s biodiversity^11^, we repeated the threat analyses on the ACBRs, here using a 25 km^2^ grid resolution because of the small size of these ecoregions. We also repeated the full set of analyses using only data from the last two decades (~ 2.54 million activity localities) to represent contemporary human activity.

**Fig. 1.**
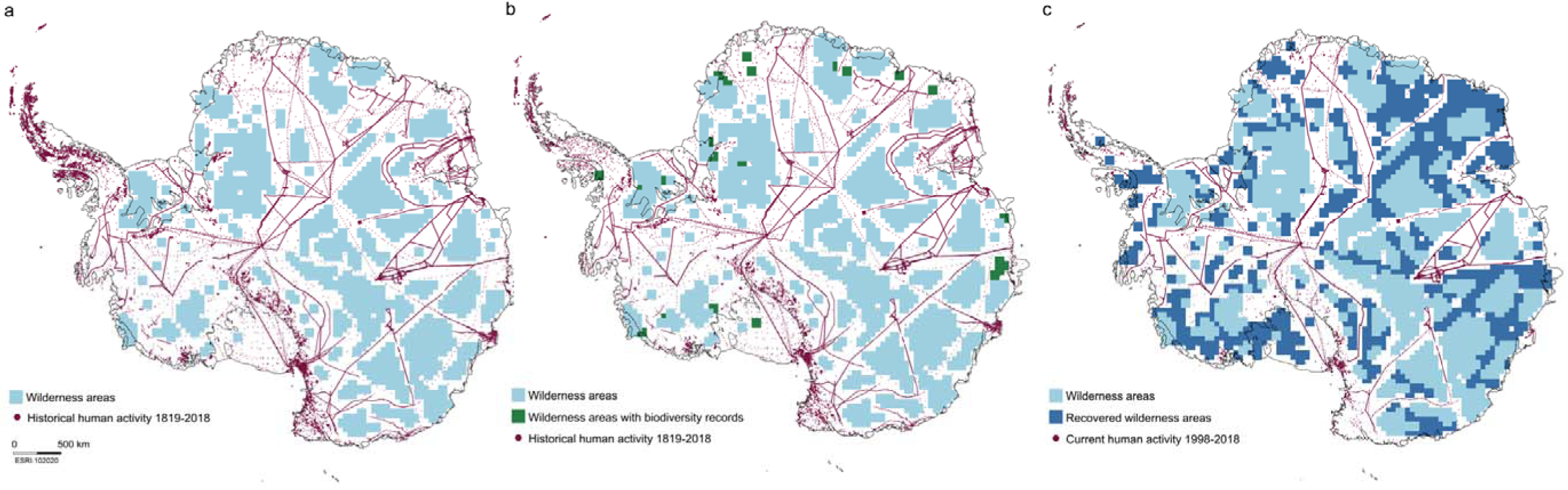
Antarctica’s wilderness and the extent of human activity. a) Wilderness areas ≥ 10 000 km^2^ (light blue squares) with no record of ground-based human activity across ~ 2.7 million activity locality records from 1819-2018 (points). b) Wilderness areas with no record of human activity, excluding biological sampling records. c) Wilderness areas with no record of ground-based human activity (points) across the complete human activity records (light blue squares) and with no records from the last twenty years (dark blue squares; 1998-2018). Many of these areas are contiguous, resulting in vast, ice-covered wilderness areas.

Antarctica’s wilderness encompasses 31.9% of the continent’s surface area (Fig. 1a). Human activity is broadly-distributed across the continent, leading to a fragmented set of wilderness areas, although large areas (up to 812 500 km^2^) of contiguous wilderness exist in East Antarctica and adjacent to the Filchner Ice Shelf. Antarctica’s wilderness does not, however, capture any sites of substantial biological value. When biological collection records were excluded from the primary data (Fig. 1b), ice-free ACBRs were largely excluded from the wilderness, with only 1006 km^2^ (1.41%) of their 71 537 km^2^ surface area represented (Fig. 2, Extended Data Table 1). None of the 220 Important Bird Areas^9^ and only 228 of the more than 48 000 species locality records in the Antarctic Terrestrial Biodiversity Database (0.47%)^12^ lie within a wilderness area. Wilderness was also entirely unrepresented in the 3809 km^2^ of Antarctic Specially Protected Areas.

**Fig. 2.**
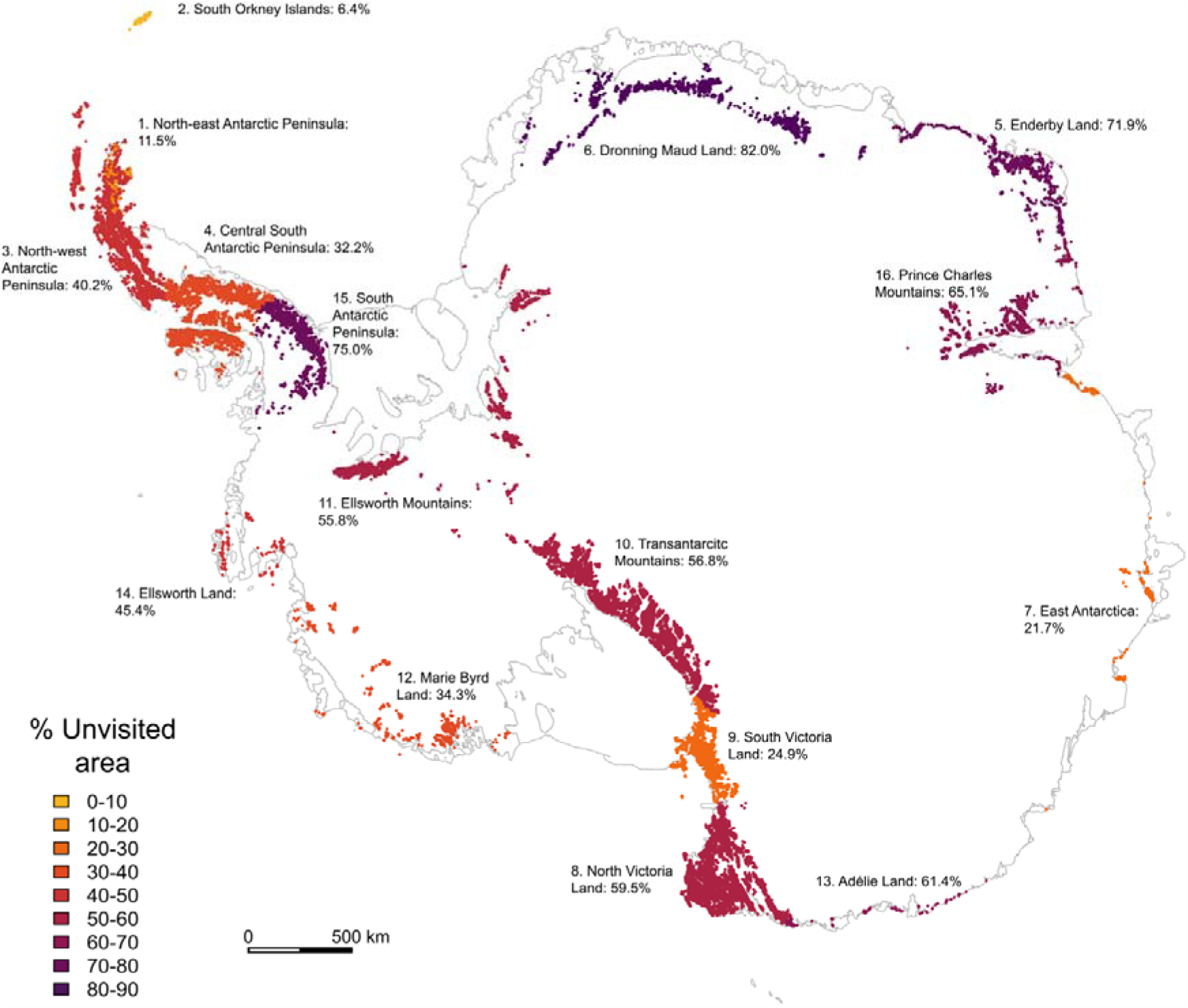
Percentage of unvisited land areas per ice-free Antarctic Conservation Biogeographic Region (ACBR), modelled at a high (25 km^2^) spatial resolution.

In terms of anthropogenic pressure, Antarctica’s wilderness areas are distant from currently-occupied research stations (Fig. 3a) and from sites of tourism landings in the 2017- 2018 season (Fig. 3b). Although the minimum distance between the edge of any single wilderness area and an occupied station was less than 2.5 km, the median distance was over 280 km (Interquartile range (IQR): 198-397 km; Extended Data Table 2). Because the majority of tourist activity is currently concentrated around the Antarctic Peninsula and McMurdo Sound regions^22^, the median distance between a wilderness area and tourist activity exceeds 990 km (IQR: 700-1408 km; Extended Data Table 2). Previous definitions of Antarctic wilderness areas have suggested that they should be at least 200 km from human activity^8^. Compared to the highly-visited regions of Antarctica, the wilderness areas are at low risk from non-native species and climate change under forecast future conditions. Only two non-native species (*Festuca rubra* and *Poa pratensis* (Poaceae)) – of the 24 found by model scenarios to be capable of establishing in Antarctica by 2100 under business as usual greenhouse gas emissions scenarios (IPCC RCP 8.5)^23^ – might be capable of establishing in 5% of the Antarctic wilderness (that is, within some portion of a 220 125 km^2^ area; Extended Data Fig. 2a). The remaining wilderness areas were climatically unsuitable for all 93 modelled non-native species, even under climate conditions forecast for 2100^23^. Because few ice-free ACBRs are contained within the wilderness, and none of these are in regions predicted to show expansion of ice-free land area under the RCP 8.5 climate scenario for 2098^11^, future ice loss will make little change to the extent to which sites typically occupied by biodiversity are included within the Antarctic wilderness.

**Fig. 3.**
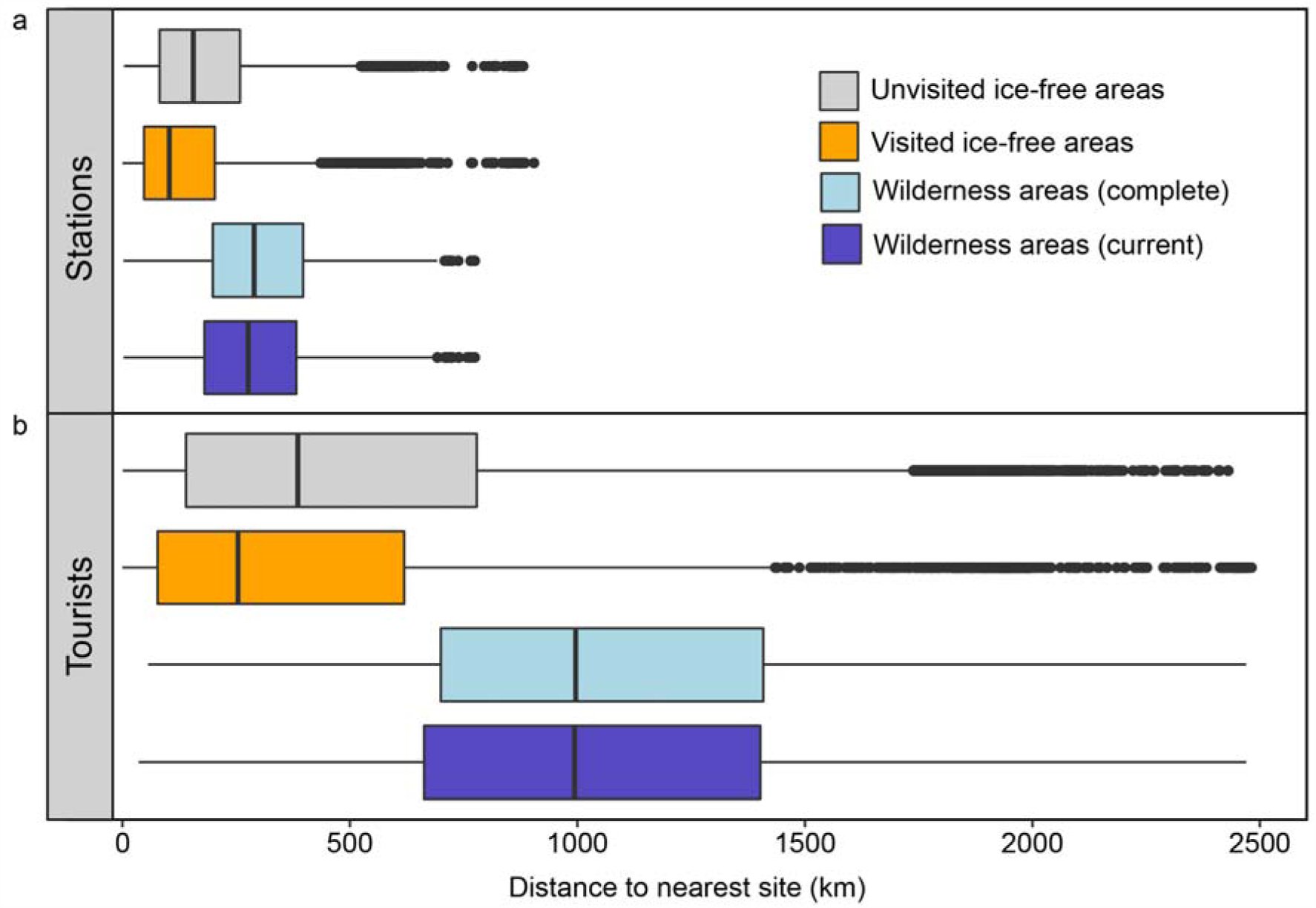
Proximity of Antarctica’s wilderness areas and ice-free areas to human activity. Minimum cartesian distances (boxes indicate median distances and interquartile range (IQR), whiskers indicate the smallest and largest observations within 1.5 × IQR, points indicate outliers) from the boundaries of wilderness areas and the centroid coordinates of ice-free areas to the nearest currently-operating research station^54^ (a), and the nearest tourist landing site from the 2017-2018 season^55^ (b). Wilderness and ice-free areas were typically more isolated from tourist activity than stations. All groups significantly different, except for the complete and current wilderness areas (see Main text).

Within the ice-free ACBRs, human activity has been extensive. At a 25 km^2^ resolution, visits have covered a mean of > 48% of the ACBR areas, although the proportion of unvisited area varied widely across the ACBRs, from less than 7% to more than 81% (Fig. 2; Extended Data Table 1). Unvisited ice-free areas were closer to research stations and sites of recent tourism landings than wilderness areas (Fig. 3; SAR stations: *z* = 17.62, P < 0.001; tourists: *z =* 2.76, P = 0.006; Extended Data Table 2-3), but were significantly more isolated from stations and tourists than visited ice-free areas (Fig. 3; SAR stations: *z* = −4.55, P < 0.0001; tourists: *z =* −5.38, P < 0.0001; Extended Data Table 2-3). For the future ice-free sites predicted to be suitable for one or more non-native species under future climate conditions (RCP 8.5 2100)^23^, there was no difference in the number of non-native species predicted to find suitable conditions across visited and unvisited ice-free sites (SAR: *z* = −0.75, P = 0.463; Extended Data Table 4). Substantial increases in ice-free area by 2098 are not expected across most of the ACBRs^11^, but in the most affected northern Antarctic Peninsula and maritime Antarctic island ACBRs (ACBRs 1-4), visited areas are expected to gain significantly more ice-free area within the next 80 years (9423 km^2^) than unvisited areas (4310 km^2^; GLMM: *t=* 6.22, df = 4, P < 0.0001; Extended Data Table 5-6).

If human activity data from only the last two decades are used, reflecting contemporary impact, the Antarctic wilderness increases from 4 357 500 km^2^ to 7 770 000 km^2^, covering 57% of the continent’s surface area (Fig. 1c). Nonetheless, it remains highly fragmented (Fig. 1c), fails to capture most of the continent’s significant biodiversity features (Supplementary File 1), and is at similar risk from anthropogenic threats (Fig. 3; Extended Data Table 2-3; Supplementary File 1).

An important caveat for our assessment is its conservatism with regard to activity data availability. Scientific data coverage favours Western countries (for example, Soviet era data are not well represented), fails to capture very recent scientific activity that has not yet resulted in data output, and omits proposed expeditions which, from their announcements, seem certain to cross areas identified here as wilderness (e.g. the planned 2018-2019 East Antarctic International Ice Sheet Traverse^27^). Moreover, although Parties to the Antarctic Treaty are expected to monitor human impacts, few do so and no centralised comprehensive repository of human activity exists^28^. Tourist data are generally better reported because they are coordinated through the International Association of Antarctica Tour Operators, but some private expedition data remain unreported, further indicating that the actual area of wilderness is probably smaller than estimated here.

Intact wilderness elsewhere on the globe is valued for its biodiversity value, including as a baseline for comparative assessments of anthropogenic impact^2,3^. Our results show that the Antarctic wilderness encompasses somewhere between a third to a half of the continent, but excludes most of its significant biodiversity. Such a situation is unique globally. Even when focusing just on Antarctica’s ice-free ecoregions (ACBRs), on average ~ 50% of their area has been subject to human activity. Nonetheless, both unvisited areas within the ACBRs and wilderness areas themselves are isolated from current anthropogenic impacts, making a strong case for their inclusion in Antarctic Specially Protected Areas. Moreover, human activity on continental Antarctica is more easily regulated than elsewhere because no indigenous populations exist, and human activity is restricted to science and tourism^24^. Evidence-based planning for and designation of new protected areas, explicitly considering the trade-offs between the benefits of science and tourism, and the importance of retaining wilderness areas or regaining their biodiversity value through the spatial restriction of human activity, could therefore readily be implemented. Such planning is well within the grasp of modern conservation science^29^. Given rapidly expanding and diversifying human activity^8,21,22^, and the general absence of Antarctica’s wilderness from its protected area system, it is also urgent. The outcomes could provide the Antarctic Treaty Parties with the mechanism required to implement their legal obligations to^13^, and renewed enthusiasm for^30^, the protection of the Antarctic environment.

## Acknowledgments

We thank K. Close, G. A. Duffy, R. Harvey, K. A. Hughes, T. Robertson and D. Smith for their assistance in identifying activity data and M. A. McGeoch, H. P. Baird and L. Chapman for reading a previous version of this manuscript. This research was supported by Australian Antarctic Science Grant 4482 and a Sir James McNeill Foundation Postgraduate Research Scholarship to R.I.L. F.M. was supported by a New Zealand Ministry of Business, Innovation and Employment grant (CO9X1413).

## Author contributions

S.L.C., B.W.T.C. and R.I.L. conceived the study. R.I.L., B.W.T.C., F.M., B.R., J.D.S. and A.T. collected the human activity data. R.I.L. conducted the analyses. R.I.L. and S.L.C. drafted the initial manuscript. All authors contributed to the final manuscript.

## Competing interests

S.L.C. is the President of the Scientific Committee on Antarctic Research.

## Additional information

**Extended data** is available for this paper at Monash Figshare (doi: 10.26180/5c32bf1b041ea).

**Supplementary data** is available for this paper at Monash Figshare (doi: 10.26180/5c32bf1b041ea).

## Reprints and permission information

### Publishers note

Springer Nature remains neutral with regard to jurisdictional claims in published maps and institutional affiliations.

## Methods

No statistical methods were used to predetermine sample size. The experiments were not randomized and the investigators were not blinded to allocation during experiments and outcome assessment.

### Historical human activity

We used publications and scientific databases to assemble an extensive record of ground-based human activity across the Antarctic continent and its offshore islands (south of 60°S) from 1819 to 2018 (see ‘Data availability’). We identified 2,698,431 activity records from 363 sources, of which 2,009,823 were unique site coordinates. Records of human activity included scientific sampling sites, traverses, infrastructure and tourism. They excluded aerial surveys, remote-sensed data and data from social networking and public image-hosting services (e.g. Twitter, Flickr, Facebook). Human activity data were available in three source formats: geographic coordinates, maps or place name records. Where available, we converted geographic coordinates from various source formats to the South Pole Lambert Azimuthal Equal Area projected coordinated system (ESRI:102020), using the sp package (ver. 1.3-1)^31^ in R (ver. 3.4.1)^32^. To extract activity data from digitized maps, we projected maps in ArcGIS (ver. 10.6)^33^, using high-resolution spatial layers of the Antarctic coastline and exposed rock outcrops, sourced from the Scientific Committee on Antarctic Research (SCAR) Antarctic Digital Database (ADD, ver. 7)^34^. Markers were then placed on map features of interest to extract activity localities.

Although place-name records varied widely in precision within and across sources (e.g. Cape Roberts hut vs. Transantarctic Mountains), they were the best-available records in several cases, such as for historical expeditions^35^. To infer the accuracy of place-name records, we sorted places by feature-type into two approximate size classes: fine-resolution features (~ ≤ 25 km^2^, e.g. Casey Station) and coarse-resolution features (> 25 km^2^, e.g. Dronning Maud Land; see Table S1 for sorting protocol). We used this resolution because it is the finest resolution we adopted for the whole-of-continent analysis. Coordinates for fine-resolution features were sourced from the SCAR Composite Gazetteer of Antarctica^36^ and included in the human activity dataset. We excluded place-name records for coarse-resolution features, such as mountain ranges, coasts and lands. Because islands and archipelagos vary greatly in size, yet have readily-delineated boundaries, the areas of visited islands were calculated to sort them into fine-resolution and coarse-resolution classes. We used the spatialEco (ver. 0.0.1-7)^37^ R-package to identify islands in the high-resolution coastline polygon of Antarctica^34^ that intersected with the SCAR Gazetteer coordinates for visited islands. The areas of visited islands were calculated using the raster (ver. 2.5-8)^38^ R-package and the SCAR Gazetteer coordinates for visited islands or island groups ≤ 25 km^2^ were included in the human activity dataset.

We sourced data from multiple databases to maximize the spatial and temporal coverage of the activity dataset. This approach likely captured duplicate data, precluding a reliable analysis of visitation frequency or impact. Even transitory visits can, however, have substantial long-term effects on Antarctic sites^26,39-43^, especially since rates of biological activity and recovery in the region are low^25,44^. Historical activity records also underrepresent human activity because some activities were undocumented, un-digitized or unpublished (e.g. exploratory field expeditions), or the visit was not captured in the literature search. Our assessment is, therefore, a conservative estimate of human visitation across Antarctica.

### High-resolution visitation grid

To identify unvisited areas in Antarctica, we used the human activity records to create a high-resolution (25 km^2^) presence/absence grid of activity across the continent, using the South Pole Lambert Azimuthal equal area projection and the R-packages raster^38^ and rgdal (ver. 1.3-4)^45^. The human activity dataset was filtered to exclude records that occurred north of 60°S or in marine areas. To exclude marine areas, a land mask of Antarctica was created by converting the high-resolution Antarctic coastline spatial polygon^34^ into a 100 × 100 m grid, where cells overlapping land areas (including ice-shelves) had a value of one, and non-land cells had a value of zero. We then aggregated this grid to the resolution of the visitation grid (25 km^2^), where cell values were calculated as the proportion cell overlap with land areas. Marine areas (i.e. those cells with no overlap with land) were assigned NA values in the human activity grid. To control for edge effects, we calculated the total amount of unvisited and visited land across Antarctica by weighting the area of each cell by the proportion of cell overlap with land using the aggregated land mask. To identify unvisited and visited sites within the ice-free Antarctic Conservation Biogeographic Regions (ACBR), we repeated this procedure using a high-resolution (30 × 30 m) grid of the ACBR spatial polygon (ver. 2)^12^, aggregated to the resolution of the visitation grid (25 km^2^), to exclude all land cells that did not overlap with ACBR areas. Because ice-free areas are typically small and highly-fragmented, the minimum size criterion for wilderness areas (≥ 10,000 km^2^) was not applied to ice-free areas.

### Identification of wilderness areas

Here, we defined Antarctic wilderness areas as all contiguous land areas ≥ 10 000 km^2^ (based on Mittermeier et al.^1^) with no record of historical human visitation. To identify wilderness areas, the high-resolution (25 km^2^) visitation grid was aggregated to a 100 × 100 km grid (= 10 000 km^2^) of Antarctica, where cells with a value of one had no visitation records and cells with a value of zero were visited (NA for marine cells). To reduce spatial bias caused by grid placement, we repeated the aggregation procedure, offset in each *x, y* spatial direction by 50 kms. The two aggregated grids were then overlaid, with the offset thus resulting in a 50 × 50 km (2 500 km^2^) grid of Antarctica used for the identification of unvisited areas. This grid was then used to identify contiguous unvisited areas of ≥ 10 000 km^2^ (i.e. ≥ four contiguous 2 500 km^2^ cells).

We created wilderness grids using i) the complete set of human activity records (1819-2018; i.e. the primary data), ii) contemporary activity records from the last twenty years (1998-2018), iii) the complete activity records excluding biological sampling data or iv) contemporary activity records from the last twenty years excluding biological sampling data. Because most data sources reported only the year or season of visitation and most Antarctic field activity occurs during the austral summer, we excluded records prior to and including the 1997-1998 field season, but included records from 1998 onwards, to quantify current human activity and additional wilderness areas which had no visitation records from the last twenty years.

Despite the increasing use of remote-sensing technologies for monitoring Antarctic biodiversity^46,47^, the vast majority of biological data have been collected from ground-observations. Likewise, bird population censuses, required to identify Important Bird Areas, are typically conducted during field expeditions^9,48^. Wilderness areas identified from a complete record of human activity will, therefore, by definition, exclude most of Antarctica’s biodiversity and IBAs. We therefore created a wilderness grid using human activity records that excluded biological sampling records to determine whether Antarctica’s wilderness areas capture the continent’s biodiversity, independent of the visitation records required to obtain the biodiversity data. Activity records from publications explicitly describing Antarctica’s biodiversity and general science records that could not be separated by discipline (e.g. Hughes et al.^41^) were excluded from the human activity records (95,986 excluded biological records, from 41 sources; see ‘Data availability’). The wilderness grids derived from the human activity data excluding biodiversity records were used to calculate the representation of biologically significant sites within Antarctica’s wilderness areas; all other analyses were conducted on the complete and contemporary wilderness grids.

### Representation of biologically significant sites

To determine whether Antarctica’s wilderness areas capture major features of biodiversity value, spatial layers of Antarctica’s ice-free Antarctic Conservation Biogeographic Regions (ACBRs)^12^, 220 Important Bird Areas (IBAs)^49^, biodiversity^12^ and Antarctic Specially Protected Areas (ASPAs)^12^ were overlaid onto the wilderness areas excluding biological sampling activity data. Ice-free areas, IBAs and species-rich sites are rare across the continent and under substantial pressure from human activity^9,21,50^, making them important conservation priorities. We used the ice-free ACBRs as a spatial layer of land areas suitable for Antarctic species because areas without the presence-only biodiversity records are not necessarily depauperate and the vast majority of Antarctic species occupy ice-free habitats^11^. We converted the wilderness grids into spatial polygon layers of the boundaries of Antarctica’s wilderness areas, using the rasterToPolygons function in the raster R-package^38^. Spatial polygons of the IBAs and ACBRs were re-projected to the same coordinate reference system as the wilderness polygons (ESRI:102020) using the rgdal R-package^45^. Representation was calculated as the area of intersection between the wilderness areas, ACBRs, IBAs, ASPAs, and the total number of biodiversity records within wilderness areas (raster R package)^38^. The biodiversity layer^12^ contains data on the number of species records per 10 × 10 km grid cell across Antarctica (total number of records = 48,795), derived from the Antarctic Terrestrial Biodiversity Database^51^ (see Terauds & Lee^12^).

### Current threats to Antarctic wilderness

Proximity to human activity has been identified as a threat to Antarctic wilderness^11,40,52^. To evaluate current threats to Antarctica’s wilderness areas, we calculated their proximity to sites of high-density human activity, including currently-operating research stations^53^ and the sites of tourism landings in the 2017-2018 season^54^. We used the wilderness areas identified using the full human activity dataset. Minimum Cartesian distances from the boundaries of wilderness areas to stations and recent tourist landing sites were calculated using the raster^38^ and rgeos R-packages (ver. 0.3-28)^55^. We also calculated the minimum distances from unvisited and visited ice-free sites in the high-resolution (25 km^2^) human activity grid to stations and tourist landing sites, using the centroid coordinates of each ice-free cell. To determine whether unvisited or visited ice-free sites were at greater risk from current human activity, we used ordinary least squares (OLS) linear regression models to determine whether visitation had an effect on the proximity of ice-free sites to stations and tourists. Distances between sites were log-transformed to improve the normality of the OLS residuals. These and other statistical tests are two-tailed. Because we found significant spatial autocorrelation in the residuals of all models, measured using Global Moran’s I (Moran’s I = 0.94-0.99, all p < 0.001), we used spatial simultaneous autoregressive (SAR) models to incorporate a spatial error term into the models to account for the spatial non-independence of the data (spdep R-package^56^, ver. 0.7-8). SAR models supplement OLS regression with a spatial weights matrix, here calculated using the four nearest neighbours of each spatial point, to reduce the amount of residual spatial autocorrelation in the models. In each case, AIC values indicated that the SAR models were a better fit to the data and had less residual spatial autocorrelation (measured using Moran’s I) than the non-spatial OLS models (Extended Data Table 3).

### Forecast threats to Antarctic wilderness

In addition to direct human activity, non-native species and climate change are the foremost threats to Antarctica’s ecosystems^11,23^. We used recent forecasts of habitat suitability for non-native species^23^ and projections of ice-melt under future climate conditions^11^ to estimate the vulnerability of Antarctica’s wilderness areas (derived from the full human activity data set), and unvisited and visited ice-free areas to future threats.

To evaluate the future climate suitability of wilderness areas for non-native species, forecast species distribution models (SDMs)^23^ were used to estimate the number of non-native species that could establish per wilderness area, and the proportion of total wilderness that might become climatically-suitable for these species by 2100. The ensemble SDMs predict the suitability of Antarctica’s land areas for 69 of the 100 world’s worst invaders and 24 cold-tolerant non-native species that could establish under the IPCC’s representative concentration pathway (RCP) 8.5 climate scenario for 2100^23^. The RCP 8.5 scenario allows for unabated emissions (“business as usual”) and is the most consistent with current observations of Antarctica’s ice losses^57^. The SDMs were re-projected to the coordinate reference system (ESRI:102020) and resolution (25 km^2^) of the human activity grid using nearest neighbour interpolation. For each spatial cell, we calculated the total number of non-native species that could establish. A spatial polygon of the wilderness areas was used to extract the maximum number of non-native species that could establish per wilderness cell and the proportion of area overlap between wilderness areas and sites predicted to be suitable for one or more species, using the raster^38^ and rgeos packages^55^ in R. We also calculated the number of non-native species that could establish per visited and unvisited ice-free cell by masking the combined SDM layer with the ice-free human activity grid (25 km^2^). We used a OLS regression to determine whether visitation had a significant effect on the number of non-native species that could establish per ice-free site for sites that were predicted to be suitable for one or more species. The number of species per site was log-transformed to improve the normality of residuals. Because we found significant spatial autocorrelation in the residuals of the model, measured using Global Moran’s I (Moran’s I = 0.97, p < 0.0001), we used a SAR model to incorporate a spatial error term into the model to account for the spatial non-independence of the data (spdep R-package^56^). The spatial weights used to estimate the error term were calculated using the four nearest neighbours of each spatial point.

To evaluate the amount of new ice-free area that is predicted to emerge within Antarctica’s wilderness areas under future climate conditions, the forecast extent of ice-free areas was extracted from Lee et al.’s^11^ mean ice-melt projections under the RCP 8.5 climate scenario for 2098. For each wilderness area, the current ice-free area was calculated from the medium-resolution rock outcrop spatial layer from the Antarctic Digital Database (ver. 7)^34^. The medium-resolution rock-outcrop layer best matches the resolution of the forecast ice-free layer^11^. We calculated the sum of the differences between the current and forecast ice-free areas as a measure of predicted environmental change within Antarctica’s wilderness areas. We also calculated the extent of ice-cover change within unvisited and visited sites per ACBR. To determine whether unvisited or visited ice-free areas are at greater risk from ice-melt under future conditions, we fitted a generalised linear mixed effect model (GLMM) with a Gamma error distribution and log link function, using the lme4 R-package (ver. 1.1-18-1)^58^, to model the effect of visitation on the total amount of ice-free area increase per ACBR, with ACBR ID included as a random effect. Because substantial ice-cover change is only predicted for the four northernmost ACBRs^11^, only data from ACBRs 1-4 were included in the model.

## Data availability

The human activity and wilderness areas spatial data have been made available through Monash Figshare (doi: 10.26180/5c32bf1b041ea).

## References

1. Mittermeier, R. A. et al. Wilderness and biodiversity conservation. Proc. Natl. Acad. Sci. USA 100, 10309–10313 (2003).

2. Watson, J. E. et al. Catastrophic declines in wilderness areas undermine global environment targets. Curr. Biol. 26, 2929–2934 (2016).

3. Watson, J. E. et al. Protect the last of the wild. Nature 563, 27–30 (2018).

4. Berkman, P. A., Lang, M. A., Walton, D. W. H. & Young, O. R. (eds.) Science Diplomacy. Antarctica, Science, and the Governance of International Spaces. (Smithsonian Institution, Washington, DC, 2011).

5. Coetzee, B. W. T., Convey, P. & Chown, S. L. Expanding the protected area network in Antarctica is urgent and readily achievable. Conserv. Lett. 10, 670–680 (2017).

6. Codling, R. in Antarctica in the Environmental Era (Dingwall, P. R. ed.) 31–37 (Department of Conservation, Wellington, New Zealand, 1998).

7. Summerson, R. in Protection of the Three Poles (Huettmann, F. ed.) 77–109 (Springer, Berlin, 2012).

8. Summerson, R. & Tin, T. Twenty years of protection of wilderness values in Antarctica. The Polar Journal, doi:10.1080/2154896x.2018.1541548 (2018).

9. Harris, C. M. et al. Important Bird Areas in Antarctica 2015. (BirdLife International and Environmental Research & Assessment Ltd., Cambridge, 2015).

10. Chown, S. L. et al. The changing form of Antarctic biodiversity. Nature 522, 431–438 (2015).

11. Lee, J. R. et al. Climate change drives expansion of Antarctic ice-free habitat. Nature 547, 49–54 (2017).

12. Terauds, A. & Lee, J. R. Antarctic biogeography revisited: updating the Antarctic Conservation Biogeographic Regions. Divers. Distrib. 22, 836–840 (2016).

13. Antarctic Treaty Consultative Meeting. Protocol on Environmental Protection to the Antarctic Treaty (Antarctic Treaty Secretariat, Buenos Aires, 1991). http://www.ats.aq/documents/recatt/Att006_e.pdf.

14. Martin, T. G. & Watson, J. E. Intact ecosystems provide best defence against climate change. Nat. Clim. Change 6, 122–124 (2016).

15. Cole, D. N. & Landres, P. B. Threats to wilderness ecosystems: impacts and research needs. Ecol. Appl. 6, 168–184 (1996).

16. Rintoul, S. R. et al. Choosing the future of Antarctica. Nature 558, 233–241 (2018).

17. Dodds, K., Hemmings, A. & Roberts, P. Handbook on the Politics of Antarctica. (Edward Elgar, London, 2017).

18. Bastmeijer, K. & Van Hengel, S. The role of the protected area concept in protecting the world’s largest natural reserve: Antarctica. Utrecht L. Rev., 5, 7–12 (2009).

19. Summerson, R & Riddle, M. J. Assessing wilderness and aesthetic values in Antarctica. In Antarctic Ecosystems. Models for Wider Ecological Understanding (pp. 303–307) (New Zealand Natural Sciences, Christchurch, 2000).

20. Chown, S.L. et al. Antarctica and the Strategic Plan for Biodiversity. PLoS Biology 15, e2001656 (2017).

21. Pertierra, L. R., Hughes, K. A., Vega, G. C. & Olalla-Tárraga, M. Á. High resolution spatial mapping of human footprint across Antarctica and its implications for the strategic conservation of avifauna. PloS One 12, e0168280 (2017).

22. Hughes, K. A. et al. Human-mediated dispersal of terrestrial species between Antarctic biogeographic regions: A preliminary risk assessment. J. Envion. Manage. 232, 73–89 (2019).

23. Duffy, G. A. Barriers to globally invasive species are weakening across the Antarctic. Divers. Distrib. 23, 982–996 (2017).

24. Tin, T., Lamers, M., Liggett, D., Maher, P. T. & Hughes, K. A. Setting the scene: human activities, environmental impacts and governance arrangements in Antarctica. In Antarctic Futures (pp. 1–24) (Springer, Dordrecht, 2014).

25. Convey, P. The influence of environmental characteristics on life history attributes of Antarctic terrestrial biota. Biological Reviews 71, 191–225 (1996).

26. Ayres, E. et al. Effects of human trampling on populations of soil fauna in the McMurdo Dry Valleys, Antarctica. Conserv. Biol. 22, 1544–1551 (2008).

27. Agence Nationale Recherche. East Antarctic International Ice Sheet Traverse (DS0101) (ANR, France) http://www.agence-nationale-recherche.fr/Project-ANR-16-CE01-0011 (2016).

28. Hughes, K. A. How committed are we to monitoring human impacts in Antarctica? Environ. Res. Lett. 5, 041001 (2010).

29. Hanson, J. O., Rhodes, J. R., Possingham, H. P., & Fuller, R. A. raptr: Representative and adequate prioritization toolkit in R. Methods in Ecology and Evolution 9, 320–330, doi:10.1111/2041-210x.12862 (2018).

30. Antarctic Treaty Consultative Meeting. Santiago Declaration (Antarctic Treaty Secretariat, Buenos Aires) https://www.ats.aq/documents/ATCM39/ad/atcm39_ad003_e.pdf (2016).

## References

31. Pebesma, E. J. & Bivand, R. S. Classes and methods for spatial data in R. R News 5, 9–13 (2005).

32. R Core Team. R: A language and environment for statistical computing. R Foundation for Statistical Computing (Vienna, Austria, 2017).

33. ESRI. ArcGIS Desktop, release 10.6 (Environmental Systems Research Institute, Redlands, CA, 2011).

34. Scientific Committee on Antarctic Research. Antarctic Digital Database, version 7. www.add.scar.org (2018).

35. Headland, R. K. Chronological List of Antarctic Expeditions and Related Historical Events (Cambridge University Press, Cambridge, 1989).

36. Scientific Committee on Antarctic Research. Composite Gazetteer of Antarctica (GCMD Metadata, 1992, updated 2014) http://gcmd.nasa.gov/records/SCAR_Gazetteer.html (2014).

37. Evans, J. S. spatialEco. R package, version 0.0.1-7 https://CRAN.R-project.org/package=spatialEco (2017).

38. Hijmans, R. J. raster: geographic data analysis and modeling. R package, version 2.6-7 https://CRAN.R-project.org/package=raster (2017).

39. Campbell, I. B., Claridge, G. G. C. & Balks, M. R. Short-and long-term impacts of human disturbances on snow-free surfaces in Antarctica. Polar Rec. 34, 15–24 (1998).

40. Hughes, K. A. & Convey, P. The protection of Antarctic terrestrial ecosystems from inter- and intra-continental transfer of non-indigenous species by human activities: a review of current systems and practices. Glob. Environ. Change 20, 96–112 (2010).

41. Hughes, K. A., Fretwell, P., Rae, J., Holmes, K. & Fleming, A. Untouched Antarctica: mapping a finite and diminishing environmental resource. Antarct. Sci. 23, 537–548 (2011).

42. O'Neill, T. A., Balks, M. R. & López-Martínez, J. Visual recovery of desert pavement surfaces following impacts from vehicle and foot traffic in the Ross Sea region of Antarctica. Antarct. Sci. 25, 514–530 (2013).

43. Tejedo, P., Pertierra, L. R. & Benayas, J. in Antarctic Futures (Tin, T., Liggett, D., Maher, P. T. & Lamers, M. eds.) 139–161 (Springer, Dordrecht, Netherlands, 2014).

44. Bargagli, R. Antarctic Ecosystems: Environmental Contamination, Climate Change, and Human Impact (Springer, Berlin, Germany, 2005).

45. Bivand, R. J., Keitt, T. & Rowlingson, B. rgdal: bindings for the 'geospatial' data abstraction library. R package, version 1.3-4. https://CRAN.R-project.org/package=rgdal (2018).

46. Casanovas, P., Black, M., Fretwell, P. & Convey, P. Mapping lichen distribution on the Antarctic Peninsula using remote sensing, lichen spectra and photographic documentation by citizen scientists. Polar Res. 34, 25633 (2015).

47. Schwaller, M. R., Lynch, H. J., Tarroux, A. & Prehn, B. A continent-wide search for Antarctic petrel breeding sites with satellite remote sensing. Remote. Sens. Environ. 210, 444–451 (2018).

48. Lynch, H. J., Naveen, R. & Fagen, W. F. Censuses of penguin, blue-eyed shag Phalacrocorax atriceps and southern giant petrel Macronectes giganteus populations on the Antarctic Peninsula, 2001-2007. Mar. Ornithol. 36, 83–97 (2008).

49. BirdLife International. Antarctic Important Bird Areas http://datazone.birdlife.org/home (Cambridge, 2018).

50. Convey, P., Hughes, K. A. & Tin, T. Continental governance and environmental management mechanisms under the Antarctic Treaty System: sufficient for the biodiversity challenges of this century? Biodiversity 13, 234–248 (2012).

51. Scientific Committee on Antarctic Research. Antarctic Biodiversity Database. https://www.scar.org/data-products/bio-database/ (2018).

52. Shaw, J. D., Terauds, A., Riddle, M. J., Possingham, H. P. & Chown, S. L. Antarctica’s protected areas are inadequate, unrepresentative, and at risk. PLoS Biol. 12, e1001888 (2014).

53. Council of Managers of National Antarctic Programs. Antarctic facilities operated by National Antarctic Programs in the Antarctic Treaty Area (South of 60° latitude South), version 3.0.1. https://www.comnap.aq, downloaded on 8 August 2018 (2018).

54. International Association of Antarctic Tour Operators. 2017–2018 Tourism statistics, http://iaato.org/tourism-statistics, downloaded on 29 October 2018 (2018).

55. Bivand, R. J. & Rundel, C. rgeos: interface to geometry engine - open source ('GEOS'). R package, version 0.3-28. https://CRAN.R-project.org/package=rgeos (2018).

56. Bivand, R. & Piras, G. Comparing implementations of estimation methods for spatial econometrics. J. Stat. Softw. 63, 1–36 (2015).

57. Slater, T. & Shepherd, A. Antarctic ice losses tracking high. Nat. Clim. Change 8, 1025–1026 (2018).

58. Bates, D., Maechler, M., Bolker, B. & Walker, S. Fitting linear mixed-effects models using lme4. J. Stat. Softw. 67, 1–48 (2015).

