## Supplementary Data for "Antarctica’s wilderness has declined to the exclusion of biodiversity"

^1^School of Biological Sciences, Monash University, Victoria 3800, Australia. ^2^Global Change Institute, University of the Witwatersrand, WITS 2050, Johannesburg, South Africa. ^3^Landcare Research New Zealand, Private Bag 92170, Auckland Mail Centre, Auckland 1142, New Zealand. ^4^Australian Antarctic Division, Department of the Environment and Energy, 203 Channel Highway, Kingston, Tasmania 7050, Australia. ^5^School of Biological Sciences, The University of Queensland, Queensland 4072, Australia.

**Extended Data**

**Extended Data Table 1 ǀ** Proportion of unvisited and visited land areas across Antarctica (including offshore islands and ice-shelves), and within each of the ice-free Antarctic Conservation Biogeographic Regions (ACBRs)^12^, modelled at a 25 km^2^ resolution. Wilderness area indicates the area of intersection between Antarctica’s wilderness areas and ACBRs.

**Extended Data Table 2 ǀ** Proximity of Antarctica’s wilderness and ice-free areas to currently-occupied research stations^53^ and sites of tourist landings in the 2017-2018 season^54^. Minimum distances (km) from the boundaries of wilderness areas (2 500 km^2^) and the centroid coordinates of ice-free areas (25 km^2^) to the nearest active station, or the nearest tourist landing site. Wilderness areas derived from complete human activity records (1819-2018) and current activity records (1998-2018).

**Extended Data Table 3 ǀ** Results from spatial simultaneous autoregressive (SAR) error models comparing the minimum distances from unvisited ice-free sites (Unvisit IF) and wilderness areas, unvisited ice-free sites and visited ice-free sites (Visit IF), and current wilderness areas derived from human activity data from the last twenty years (1998-2018; Wild_current_) and complete activity wilderness areas (1819-2018; Wild_comp._), to the nearest currently-operating research station^53^, or the nearest site of tourist landings from the 2017-2018 season^54^. Minimum distances between sites were log-transformed in the analysis.

**Extended Data Table 4 ǀ** Total number of unvisited and visited ice-free cells (25 km^2^) suitable for at least *n* non-native species under forecast business as usual climate conditions (RCP 8.5) for 2100^23^. Mean number of species across all sites and sites that are expected to be climatically suitable for at least one non-native species in the future.

**Extended Data Table 5 ǀ** Area of current and forecast (2098 RCP 8.5 conditions)^11^ ice-free areas at sites with no human activity records (unvisited) and visited sites within the 16 ice-free Antarctic Conservation Biogeographic Regions (ACBRs)^12^.

**Extended Data Table 6 |** Generalised linear mixed effect model (Gamma error distribution, log link function) outcomes for the effect of visitation on the total amount of ice-free area increase predicted under forecast climate conditions (RCP 8.5)^11^ for 2098 per ice-free Antarctic Conservation Biogeographic Region (ACBR)^12^, for the four northernmost ACBRs (ACBRs 1-4). Visited sites are expected to gain significantly more ice-free area within the next 80 years than unvisited ice-free areas.

**Extended Data Fig. 1 ǀ** Number of historical human activity records per 50 x 50 km cell across Antarctica, from 1819-2018 (~2.7 million records). Dark purple lines indicate the routes of recent overland traverses (e.g. 2007-2008 Norwegian-U.S. Scientific Traverse of East Antarctica), where geo-positioning data were collected automatically at a high temporal resolution (~10 minutes), resulting in many records for relatively transitory site visits.

**Extended Data Fig. 2 ǀ** Maximum number of terrestrial non-native species from the 93 species modelled by Duffy et al.^23^ for which the climate of some portion of Antarctica’s wilderness areas, derived from the complete human activity records (a; 1819-2018) and current activity records (b; 1998-2018), were predicted to be suitable under averaged, business as usual (RCP 8.5) climate conditions by 2100. Total wilderness area for which some portion is forecast to be suitable for ≥ 1 non-native species is 220 125 km^2^ for complete wilderness areas (a) and 466 025 km^2^ for current wilderness areas (b).

**Extended Data Table 1 ǀ** Proportion of unvisited and visited land areas across Antarctica (including offshore islands and ice-shelves), and within each of the ice-free Antarctic Conservation Biogeographic Regions (ACBRs)^12^, modelled at a 25 km^2^ resolution. Wilderness area indicates the area of intersection between Antarctica’s wilderness areas and ACBRs. Δ

| ACBR ID | ACBR Name | Total area (km^2^) | Unvisited area (km^2^) | Visited area (km^2^) | % area unvisited | Wilderness  area (km^2^) |
| --- | --- | --- | --- | --- | --- | --- |
| 1 | North-east Antarctic Pen. | 1221 | 141 | 1080 | 11.52 | 0 |
| 2 | South Orkney Islands | 160 | 10 | 150 | 6.40 | 0 |
| 3 | North-west Antarctic Pen. | 5241 | 2109 | 3132 | 40.23 | 0 |
| 4 | Central south Antarctic Pen. | 5004 | 1610 | 3394 | 32.18 | 0 |
| 5 | Enderby Land | 2174 | 1563 | 610 | 71.92 | 0 |
| 6 | Dronning Maud Land | 5566 | 4562 | 1004 | 81.96 | 153 |
| 7 | East Antarctica | 1089 | 236 | 853 | 21.69 | 0 |
| 8 | North Victoria Land | 9481 | 5641 | 3839 | 59.50 | 0 |
| 9 | South Victoria Land | 10238 | 2547 | 7690 | 24.88 | 0 |
| 10 | Transantarctic Mountains | 18594 | 10563 | 8032 | 56.81 | 47 |
| 11 | Ellsworth Mountains | 2872 | 1603 | 1269 | 55.81 | 0 |
| 12 | Marie Byrd Land | 1142 | 392 | 750 | 34.29 | 0 |
| 13 | Adélie Land | 171 | 105 | 66 | 61.35 | 0 |
| 14 | Ellsworth Land | 220 | 100 | 120 | 45.38 | 0 |
| 15 | South Antarctic Pen. | 2883 | 2162 | 720 | 75.01 | 0 |
| 16 | Prince Charles Mountains | 6054 | 3939 | 2115 | 65.06 | 83 |
| Total | Ice-free areas | 72 108 | 37 282 | 34 826 | 51.70 | 283 |
|  | All land areas (in. ice-shelves) | 13 650 189 | 13 057 731 | 592 458 | 95.66 | 4 357 500 |

Δ Because of resolution and rounding errors, total ACBR areas differ slightly from those published in Terauds & Lee^12^ and the % area unvisited differs slightly from the unvisited area divided by the total ACBR area in some instances.

**Extended Data Table 2 ǀ** Proximity of Antarctica’s wilderness and ice-free areas to currently-occupied research stations^53^ and sites of tourist landings in the 2017-2018 season^54^. Minimum distances (km) from the boundaries of wilderness areas (2 500 km^2^) and the centroid coordinates of ice-free areas (25 km^2^) to the nearest active station, or the nearest tourist landing site. Wilderness areas derived from complete human activity records (1819-2018) and current activity records (1998-2018).

|  |  | Minimum | Median | Mean (± SD) | Maximum |
| --- | --- | --- | --- | --- | --- |
| Wilderness Areas |  |  |  |  |  |
| Complete | Stations | 2.42 | 289.45 | 304.35 (140.86) | 774.62 |
|  | Tourists | 56.31 | 996.51 | 1042.52 (493.80) | 2469.84 |
| Current | Stations | 2.42 | 276.43 | 289.03 (141.60) | 774.62 |
|  | Tourists | 35.59 | 993.90 | 1038.98 (494.97) | 2469.84 |
| Ice-free areas |  |  |  |  |  |
| Unvisited sites | Stations | 3.6 | 155.74 | 180.43 (125.33) | 881.26 |
|  | Tourists | 2.89 | 385.90 | 562.46 (548.28) | 2430.57 |
| Visited sites | Stations | 0.24 | 103.07 | 148.05 (141.04) | 904.65 |
|  | Tourists | 0.09 | 253.98 | 445.87 (520.37) | 2481.70 |

**Extended Data Table 3 ǀ** Results from spatial simultaneous autoregressive (SAR) error models comparing the minimum distances from unvisited ice-free sites (Unvisit IF) and wilderness areas, unvisited ice-free sites and visited ice-free sites (Visit IF), and current wilderness areas derived from human activity data from the last twenty years (1998-2018; Wild_current_) and complete activity wilderness areas (1819-2018; Wild_comp._), to the nearest currently-operating research station^53^, or the nearest site of tourist landings from the 2017-2018 season^54^. Minimum distances between sites were log-transformed in the analysis.

|  |  | Coefficient | SE | *z* | λ | *n* | *P* | *I*_OLS_ (SD) | *I*_SAR_ (SD) | AIC_OLS_ | AIC_SAR_ |
| --- | --- | --- | --- | --- | --- | --- | --- | --- | --- | --- | --- |
| **Stations** |  |  |  |  |  |  |  |  |  |  |  |
| Unvisit IF | Wild_comp._ | 0.17 | 0.01 | 17.62 | 0.98 | 14397 | < 0.001 | 0.98 (174.86) | 0.08 (14.87) | 34716 | -18936 |
| Unvisit IF | Visit IF | -0.008 | 0.002 | -4.55 | 0.98 | 20623 | < 0.001 | 0.96 (203.71) | 0.05 (10.08) | 56949 | -21837 |
| Wild_current_ | Wild_comp._ | < 0.01 | < 0.01 | 1.44 | 0.97 | 4851 | 0.150 | 0.94 (100.69) | 0.12 (13.1) | 8557.8 | -5343.4 |
| **Tourists** |  |  |  |  |  |  |  |  |  |  |  |
| Unvisit IF | Wild_comp._ | 0.02 | 0.01 | 2.76 | 0.99 | 14397 | 0.006 | 0.99 (176.87) | 0.04 (7.29) | 46155 | -25754 |
| Unvisit IF | Visit IF | -0.01 | 0.002 | -5.38 | 0.98 | 20623 | < 0.001 | 0.98 (206.54) | -0.04 (-7.93) | 72355 | -18411 |
| Wild_current_ | Wild_compl._ | < 0.01 | < 0.01 | -0.38 | 0.99 | 4851 | 0.705 | 0.99 (107.18) | 0.31 (33.60) | 8675.4 | -18670 |

*I*_OLS_ (SD): Global Moran’s I statistic and standard deviation for the non-spatial Ordinary Least Squares (OLS) regression model. *I*_SAR_ (SD): Global Moran’s I and standard deviation for the spatial SAR model. In all cases, the SAR model had less residual spatial autocorrelation (lower Moran’s I) and greater goodness-of-fit (lower AIC) than the non-spatial OLS model.

**Extended Data Table 4 ǀ** Total number of unvisited and visited ice-free cells (25 km^2^) suitable for at least *n* non-native species under forecast business as usual climate conditions (RCP 8.5) for 2100^23^. Mean number of species across all sites and sites that are expected to be climatically suitable for at least one non-native species in the future.

| Species (*n*) | Unvisited | Visited |
| --- | --- | --- |
| 0 | 10514 | 6748 |
| 1 | 1432 | 578 |
| 2 | 68 | 30 |
| 3 | 25 | 13 |
| 4 | 19 | 15 |
| 5 | 45 | 43 |
| 6 | 8 | 2 |
| 7 | 36 | 34 |
| 8 | 19 | 46 |
| 9 | 64 | 26 |
| 10 | 107 | 63 |
| 11 | 27 | 20 |
| 12 | 15 | 42 |
| 13 | 13 | 9 |
| 14 | 114 | 45 |
| 15 | 15 | 32 |
| 16 | 12 | 0 |
| 17 | 24 | 12 |
| 18 | 41 | 87 |
| 19 | 38 | 87 |
| 20 | 11 | 12 |
| 21 | 3 | 10 |
| 22 | 4 | 15 |
| Mean (± SD) number species (all sites) | 0.71 (± 2.73) | 1.08 (± 3.74) |
| Mean (± SD) number species (sites ≥ 1 species) | 4.18 (± 5.43) | 7.03 (± 7.03) |

**Extended Data Table 5 ǀ** Area of current and forecast (2098 RCP 8.5 conditions)^11^ ice-free areas at sites with no human activity records (unvisited) and visited sites within the 16 ice-free Antarctic Conservation Biogeographic Regions (ACBRs)^12^. Δ

| ACBR ID | ACBR Name | Total area (km^2^) | Current unvisited area (km^2^) | Forecast unvisited area (km^2^) | Forecast increase in ice-free area (km^2^) at unvisited sites | Current visited area (km^2^) | Forecast visited area (km^2^) | Forecast increase in ice-free area (km^2^) at visited sites |
| --- | --- | --- | --- | --- | --- | --- | --- | --- |
| 1 | North-east Antarctic Pen. | 1312 | 138 | 347 | 209 | 1174 | 2480 | 1306 |
| 2 | South Orkney Islands | 160 | 10 | 91 | 81 | 150 | 506 | 356 |
| 3 | North-west Antarctic Pen. | 5432 | 2079 | 5706 | 3627 | 3353 | 10168 | 6815 |
| 4 | Central south Antarctic Pen. | 4891 | 1378 | 1758 | 380 | 3513 | 4406 | 894 |
| 5 | Enderby Land | 2134 | 1470 | 1470 | 0 | 664 | 668 | 5 |
| 6 | Dronning Maud Land | 5469 | 4452 | 4452 | 0 | 1017 | 1017 | 0 |
| 7 | East Antarctica | 1059 | 184 | 189 | 5 | 875 | 913 | 37 |
| 8 | North Victoria Land | 9397 | 5441 | 5445 | 4 | 3956 | 3959 | 3 |
| 9 | South Victoria Land | 9909 | 2048 | 2048 | 0 | 7862 | 7862 | 0 |
| 10 | Transantarctic Mountains | 18352 | 9830 | 9830 | 0 | 8522 | 8522 | 0 |
| 11 | Ellsworth Mountains | 2848 | 1507 | 1507 | 0 | 1340 | 1340 | 0 |
| 12 | Marie Byrd Land | 1097 | 329 | 331 | 2 | 768 | 771 | 4 |
| 13 | Adélie Land | 167 | 100 | 100 | 0 | 67 | 67 | 0 |
| 14 | Ellsworth Land | 214 | 92 | 92 | 0 | 121 | 123 | 2 |
| 15 | South Antarctic Pen. | 2877 | 2137 | 2137 | 0 | 740 | 740 | 0 |
| 16 | Prince Charles Mountains | 5962 | 3816 | 3818 | 2 | 2147 | 2148 | 1 |
|  | Total increase in ice-free area: | | | | 4310.28 |  |  | 9422.55 |

Δ Small differences in the current ACBR areas between Extended Data Tables 1 and 5 are a consequence of using a coarser-resolution ice-free area spatial polygon for the forecast ice-free area change analysis because this resolution best matches that used to develop the forecast models^11^.

**Extended Data Table 6 |** Generalised linear mixed effect model (Gamma error distribution, log link function) outcomes for the effect of visitation on the total amount of ice-free area increase predicted under forecast climate conditions (RCP 8.5)^11^ for 2098 per ice-free Antarctic Conservation Biogeographic Region (ACBR)^12^, for the four northernmost ACBRs (ACBRs 1-4). Visited sites are expected to gain significantly more ice-free area within the next 80 years than unvisited ice-free areas.

| Fixed effect | Coefficient | SE | 95% CI | *t* | df | *P* |
| --- | --- | --- | --- | --- | --- | --- |
| Visitation (visited sites) | 1.19 | 0.19 | 0.82-1.57 | 6.22 | 4 | 5.14e-10 |
| Random effect | Variance | SD |  |  |  |  |
| ACBR ID | 0.97 | 0.99 |  |  |  |  |

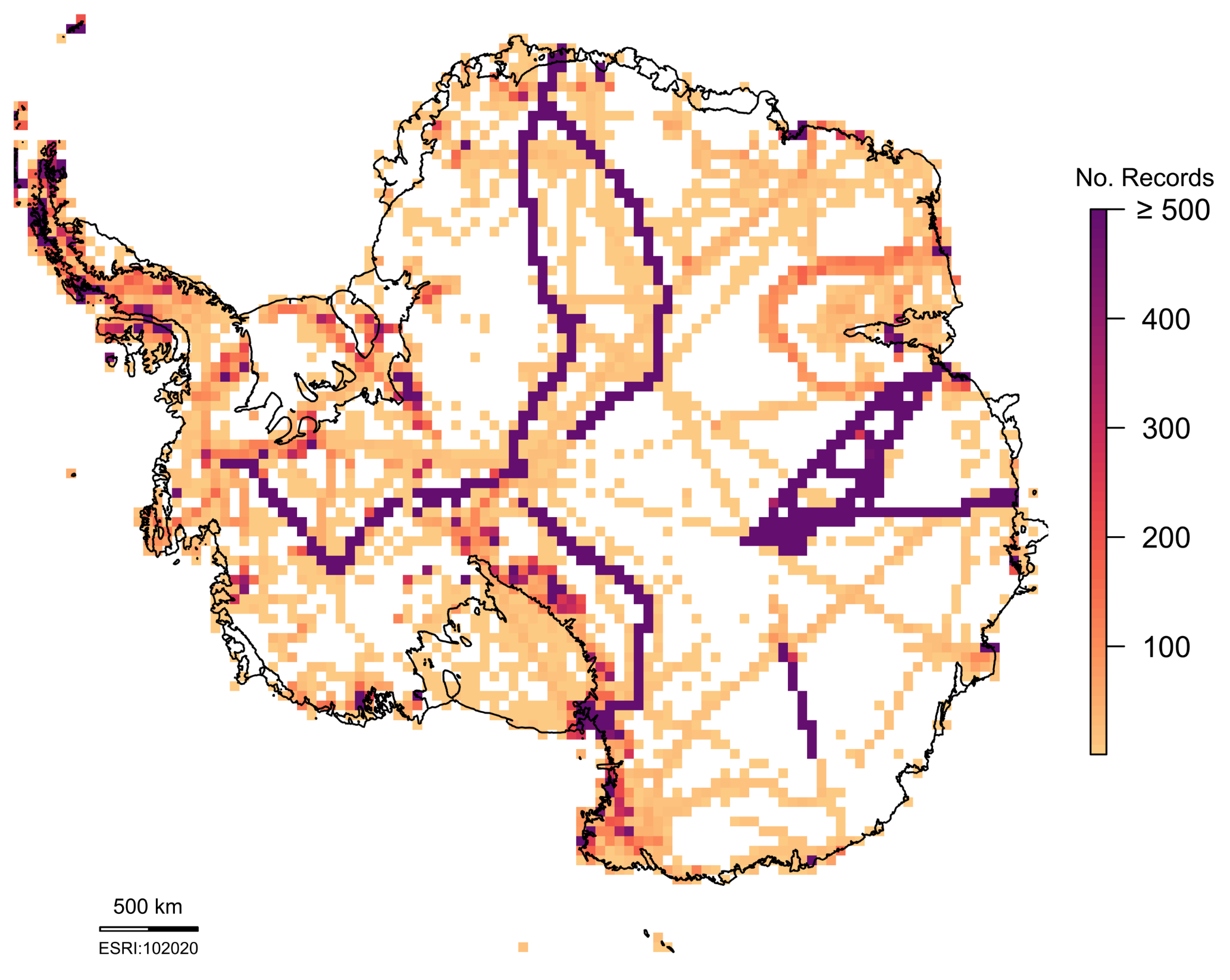

**Extended Data Fig. 1 ǀ** Number of historical human activity records per 50 x 50 km cell across Antarctica, from 1819-2018 (~2.7 million records). Dark purple lines indicate the routes of recent overland traverses (e.g. 2007-2008 Norwegian-U.S. Scientific Traverse of East Antarctica), where geo-positioning data were collected automatically at a high temporal resolution (~10 minutes), resulting in many records for relatively transitory site visits.

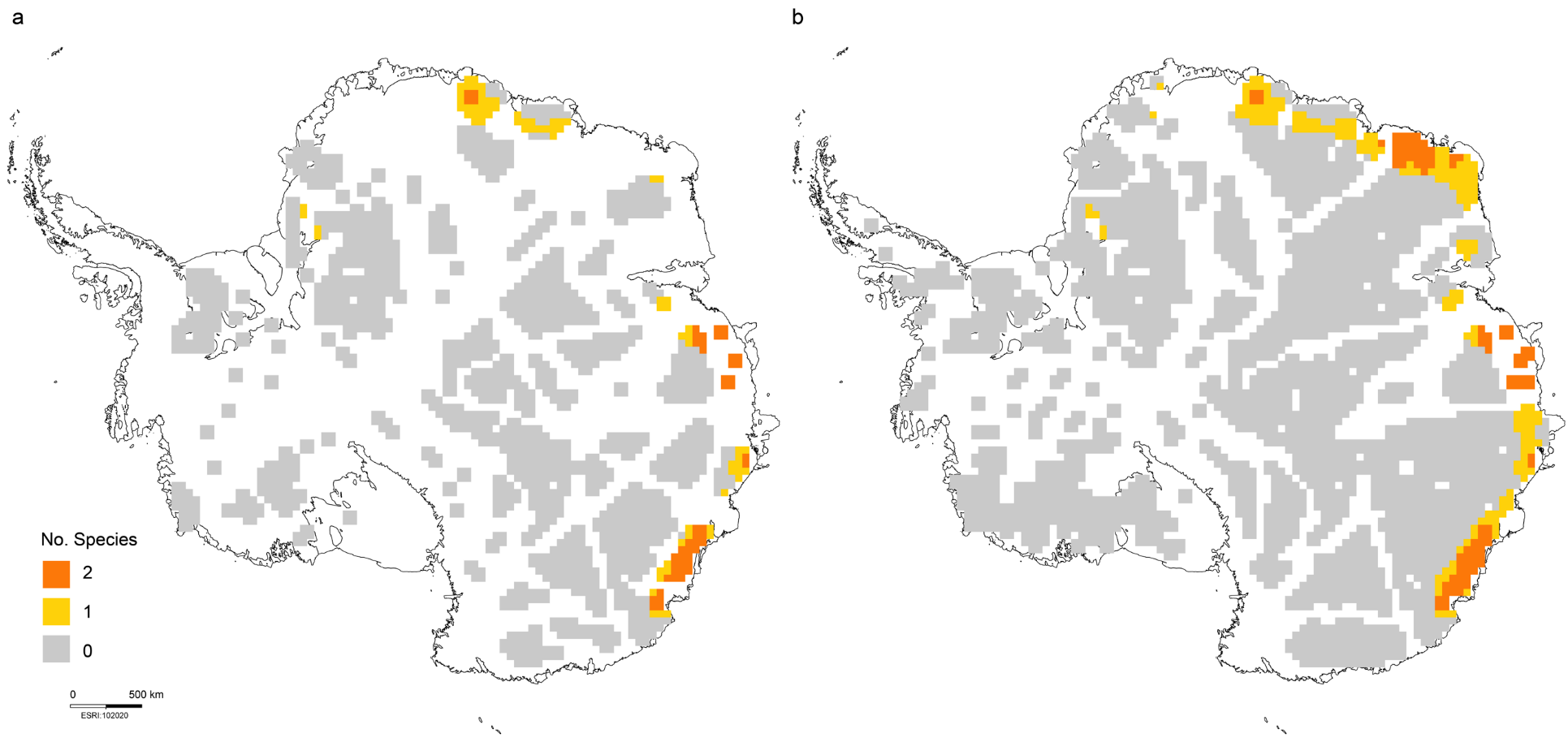

**Extended Data Fig. 2 ǀ** Maximum number of terrestrial non-native species from the 93 species modelled by Duffy et al.^23^ for which the climate of some portion of Antarctica’s wilderness areas, derived from the complete human activity records (a; 1819-2018) and current activity records (b; 1998-2018), were predicted to be suitable under averaged, business as usual (RCP 8.5) climate conditions by 2100. Total wilderness area for which some portion is forecast to be suitable for ≥ 1 non-native species is 220 125 km^2^ for complete wilderness areas (a) and 466 025 km^2^ for current wilderness areas (b).

**Supplementary Files**

**Supplementary File 1 ǀ Current wilderness**

Using only human activity records from the last twenty years (1998-2018; > 2.5 million records, of which > 1.9 million are unique localities), Antarctica’s current wilderness covers 56.92% of the continent (7 770 000 km^2^; Fig. 1c). In accordance with the wilderness derived from the full human activity dataset, when biological collection records were excluded from the current human activity records, the constrained current wilderness areas capture few features of biodiversity value. Ice-free ACBRs were largely excluded from the current wilderness, with only 11 829 km^2^ of their 71 537 km^2^ surface area (16.53%) represented. Only one of the 220 Important Bird Areas^1^ (IBA) (Sims Island; 70 ha) and 1955 biodiversity records of the more than 48 000 records in the Antarctic Terrestrial Biodiversity Database^2^ (4.0%) lie within a current wilderness area. We found no record of visitation to the Sims Island IBA since 1996^3^, while more recent investigations of the island’s bird colonies have been made from overflights and satellite imagery^4,5^. Some portion of one of the Antarctic Specially Protected Areas^2^ overlaps with the current wilderness (72.2 km^2^ of the Ablation Valley and Ganymede Heights ASPA (No. 147; Alexander Island), out of the 3809 km^2^ of total ASPAs (1.9%). Activity in this ASPA is mostly concentrated around the lakes and camp sites along its eastern boundary and no record of visitation was captured for the western sites in the human activity data records from 1998-2018 that excluded biological collection records and general science records that could not be separated by discipline.

Antarctica’s current wilderness areas (derived from the full human activity dataset from 1998-2018) are under similar anthropogenic pressure to the complete wilderness, with no significant difference between their proximity to scientific stations or sites of tourist landings in the 2017-2018 season (Fig. 3; Extended Data Table 2-3). Current wilderness areas are at low risk from non-native species and climate change, compared to the northern Antarctic Peninsula and its offshore islands^6,7^. In accordance with the complete wilderness analysis, only two non-native species (*Festuca rubra* and *Poa pratensis* (Poaceae)) of the 93 species modelled by Duffy et al.^6^ were found to be capable of establishing within current wilderness areas by 2100 under the business as usual (IPCC RCP 8.5) climate scenario (Extended Data Fig. 1b). A greater proportion of the current wilderness than the complete wilderness was predicted to be suitable for at least one non-native species in the future (Extended Data Fig. 2; 6.00% of the current wilderness (466 025 km^2^), 5.05% of the wilderness using all activity data (220 125 km^2^)). Because Antarctica’s current wilderness areas largely exclude the regions of Antarctica predicted to be most vulnerable to ice-free area expansion under future conditions^7^, the predicted increase in ice-free wilderness area under a business as usual (RCP 8.5) climate scenario by 2098 was 34.93 km^2^.

1. BirdLife International. Antarctic Important Bird Areas http://datazone.birdlife.org/home (Cambridge, 2018).
2. Terauds, A. & Lee, J. R. Antarctic biogeography revisited: updating the Antarctic Conservation Biogeographic Regions. *Divers. Distrib.* **22**, 836-840 (2016).
3. Hathway, B. Sims Island: first data from a Pliocene alkaline volcanic centre in eastern Ellsworth Land. *Antarct. Sci.* **13**, 87-88 (2001).
4. Convey, P., Hopkins, D. W., Roberts, S. J. & Tyler, A. N. Global southern limit of flowering plants and moss peat accumulation. *Polar Res.* **30**, 8929 (2011).
5. Lynch, H. J. & LaRue, M. A. First global census of the Adélie Penguin. *The Auk* **131**, 457-466 (2014).
6. Duffy, G. A. Barriers to globally invasive species are weakening across the Antarctic. *Divers. Distrib.* **23**, 982-996 (2017).
7. Lee, J. R. et al. Climate change drives expansion of Antarctic ice-free habitat. *Nature* **547**, 49-54 (2017).

**Table S1 ǀ** Initial filter used to sort place-name activity records into fine-resolution (≤ 25 km^2^) and coarse-resolution (> 25 km^2^) geographic feature classes. Islands and island groups were sorted independently by their size (see Methods). Δ

| **Fine-resolution features** | | **Coarse-resolution features** | |
| --- | --- | --- | --- |
| Airbase | Massif | Archipelago | Oasis |
| Airfield | Monument | Bay | Passage |
| Airway | Monolith | Channel | Peaks |
| Anchorage | Mount | Coast | Peninsula |
| AWS | Nunatuk | Dome | Plain |
| Base | Observatory | Fjella | Promontory |
| Beach | Point | Fjord | Range |
| Bellows | Peak | Glacier | Ranges |
| Bluff | Pinnacles | Gulf | Region |
| Camp | Pointe | Harbor | Sea |
| Cape | Pole | Harbour | Sound |
| Cemetery | Port | Hills | Strait |
| Cliffs | Reef | Ice stream | Terre |
| Cove | Reefs | Ice Tongue | Valley |
| Field | Refuge | Ice-shelf | Valleys |
| Grave | Ridge | Icesheet |  |
| Head | Rock | Island |  |
| Hill | Rocks | Islands |  |
| Hut | Rookery | Kyst |  |
| Iceport | Runway | Land |  |
| Inlet | Skiway | Plateau |  |
| Island | Station | Mountains |  |
| Lake |  | Nunatuks |  |

Δ Where small differences in spelling or use were present, such as ‘ice shelf’ and ‘ice-shelf’, these were made consistent to a single use prior to analysis.
